# Yellow Fever, Dengue Fever and West Nile Viruses Co-Circulation in Ogbomoso

**DOI:** 10.1101/265819

**Authors:** E.K. Oladipo, E.H. Awoyelu, J.K. Oloke

## Abstract

Arboviruses are an emerging threat of significant impact on human health and well-being. With increasing proportion of the world living in urban environments, inadvertently, there is the creation of better habitats for vector species. This study is aimed at establishing the occurrence of arboviruses within Ogbomoso, with a view to providing baseline data for further study. Ninety-three plasma samples from consenting individuals in the age range 1-75 years were collected and screened for IgM to dengue fever (DENV), West Nile (WNV) and yellow fever (YFV) viruses using third generation Enzyme Linked Immuno-Sorbent Assay (WKEA Med Supplies Corp, China) kits. An overall prevalence of 52.7% (49/93) were recorded from the recruited individuals. IgM antibodies to Dengue, Yellow fever and West Nile viruses were found in 16/49 (17.2%), 16/49 (17.2%) and 17/49 (18.3%), respectively. High prevalence were recorded in the age groups 16-30, 31-45 and 61-75 years. Gender analysis of the positive samples showed higher prevalence among females than males. The result also showed high prevalence in urban settings than rural settings for DENV and WNV, however, for YNF, higher prevalence was found in the rural area. The prevalence of dual and trio arboviral infection showed 17.2% and 11.8% respectively. This study confirms the circulation of Dengue fever, Yellow fever and West Nile viruses in Ogbomoso and therefore suggest the need for public awareness on vector control.

## Introduction

Arboviruses (arthropod-borne viruses), belonging to the genus *Flavivirus*, family *Flaviviridae*, are transmitted to humans primarily through the bites of mosquitoes and ticks (Lindsey *et al.,* 2014). With regards to disease impact, Flaviviruses include dengue virus (DENV), yellow fever virus (YFV), West Nile virus (WNV), etc. (Chisom *et al*., 2017). Depending on specific vectors and different natural hosts, flaviviruses have distinct geographical distributions. There is heavy build-up of mosquitoes in the tropics (Adeleke *et al.,* 2010).

Several factors that favour the spread and transmission of arboviruses include human movement, climatic and environmental factors. Climatic factors, most especially temperature, relative humidity and rainfall patterns are strong environmental drivers of vector-borne disease transmission which in turn determines the environmental suitability or potential for viral transmission (Kanfui, 2017). Activities in the urban and rural areas are of particular concern in the transmission of mosquito-borne diseases (Monath, 1994). With the relatively recent long distance travel, the spread of invasive species, particularly arboviruses, have only increased in frequency and severity (Adeleke *et al*., 2010).

DENV is known to be the most rapidly spreading mosquito-borne viral disease in the world. In the last 50 years, its incidence has increased 30-fold with increasing geographic expansion to new countries and from urban to rural settings (WHO, 2009). An estimated 50 million dengue infections occur annually with over 2.5 billion people living in dengue endemic countries (WHO, 2008). In recent years, dengue fever has been documented in travelers returning from several West African countries (Franco *et al.,* 2010). In Nigeria, DENV IgM seroprevalences of 30.8% (Faneye *et* al., 2013) and 17.2% (Oladipo *et* al., 2014) were reported. Further study showed prevalence rates of 35% and 73% (Oyero and Ayukekbong, 2014.

Secondly, twenty five countries within the sub-Saharan region are now endemic to yellow fever virus infection (Jentes *et al.,* 2011). In the last decade, the number of West African countries reporting yellow fever to World Health Organization (WHO) has increased, with 93% of the countries notifying cases in the past 4 years. This is a 30% increase compared to the period between 1995 and 1999 (WHO, 2014).

Thirdly, West Nile virus (WNV) infection has been reported in USA, Europe and few countries in Africa (Karabatos, 1985; Peiris and Amerasinghe, 1994; Hubalek and Halouzka, 1996). In 2013, a total of 2,469 WNV cases including 1,267 (51%) neauroinvasive cases were reported from 725 counties in 47 states in the USA alone (Lindsey *et al.,* 2014). Few data on WNV infection in sub-Saharan Africa is due to lack of sustained surveillance activities (Cabre *et al.,* 2006).

These data are consistent with the fact that Flaviviruses are endemic and emerging infection in Nigeria. However, the diseases are neglected, under recognized and under reported in Nigeria, particularly Ogbomoso. Hence, this study was designed to identify some arboviral causes of fever in patients attending hospitals in a rural and semi-urban center in sub-Saharan African country. This is in order to ascertain the current epidemiological situation and provide information to clinicians to enhance patient care.

## Materials and Methods

### Study Design

Ninety-three plasma samples were tested from people between 1-75 years of age in Ayedaade (urban) area of Ogbomoso North Local Government area (LGA) and Otamokun village of Ogo Oluwa LGA in Oyo State. Recruited individuals enrolled from urban and rural areas were 47 (50.5%) and 46 (49.5%) respectively. There were 56 females and 37 males. Of this, 24 (25.8%) were single and 69 (74.2%) married.

### Laboratory diagnosis

Commercial IgM ELISA kits for Dengue, West Nile and Yellow Fever were obtained from WKEA MED SUPPLIES CORP (Changchun, China) and stored at 2-8°C. Assay procedure was carried out according to the manufacturer’s specifications. Plates were labeled appropriately leaving two wells each on the last column for positive control, negative control and blank. 40µl of sample diluents was then added to 10µl of sample in each labeled well. It was carefully mixed and incubated at 37°C for 30mins. Wash buffer was prepared by diluting it 50 times before washing five times using a plate washer (Biotek^@^USA). 50 µl of enzyme conjugate was then added to all wells except the blank well and incubated at 37°C for 30mins as described above. Another round of washing was done for five times. 50µl each of chromogenic substrate A and B were then added and incubated in a dark room at 37°C for 15mins. 50µl of stop solution was then added to each well and a colour change was observed from blue to yellow. Optical density was measured using a spectrophotometer with wavelength set at 450nm. The presence or absence of dengue, West Nile and yellow fever virus IgM was determined by comparing the sample absorbance with the absorbance of the cut-off calibrator.

## Statistical analysis

Information from questionnaire was entered into an excel spreadsheet and analyzed using SPSS 16.0.

## Results

Forty-nine (52.7%) of the 97 sera were positive for DENV, WNV and YFV infections. Out of the 49 (52.7%) IgM positive patients, 16 (17.2%) were positive for DENV, 17 (18.3%) for WNV and 16 (17.2%) for YFV (**Fig. 1**). Gender analysis of the positive samples showed higher prevalence among females than males (**Table 1**). High prevalence were recorded in the age groups 16-30, 31-45 and 61-75 years in the age distribution of IgM positivity (**Table 2**). The result also shows that patients from the urban settings had more of the infections than those in the rural areas, however, for YNF, higher prevalence was found in the rural area (**Fig. 2**). **Fig. 3** shows the prevalence of dual and trio arboviral infection.

**Fig 1.**
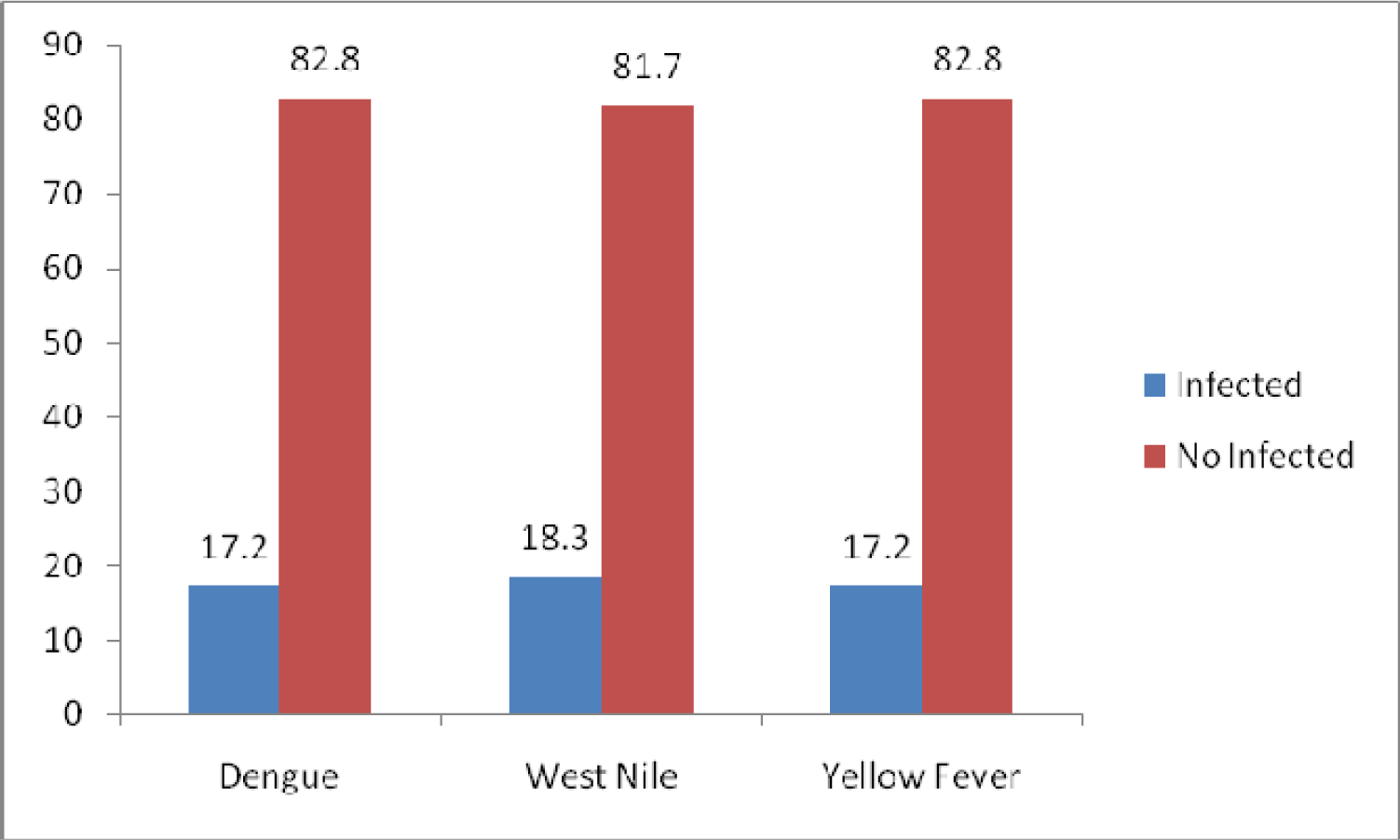
Distribution of individuals infected by DENV, WNV and YFV in the general population.

**Table 1.** Sex distribution of IgM positive individuals.

| Sex/Infection | DENV IgM+ | WNV IgM+ | YFV IgM+ |
| --- | --- | --- | --- |
| **MALE** | 7 (43.8%) | 7 (41.2%) | 6 (37.5%) |
| **FEMALE** | 9 (56.2%) | 10 (58.8%) | 10 (62.5%) |

**Table 2.** Age distribution of IgM positive people.

| Age (yrs) | DENV IgM+ | WNV IgM+ | YFV IgM+ |
| --- | --- | --- | --- |
| 1 – 15 | 2 (12.5%) | 2 (11.8%) | 2 (12.5%) |
| 16 – 30 | 4 (25.0%) | 5 (29.4%) | 4 (25.0%) |
| 31 – 45 | 6 (6.5%) | 5 (29.4%) | 5 (31.3%) |
| 46 – 60 | 3 (18.8%) | 4 (23.5%) | 1 (6.3%) |
| 61 – 75 | 1(6.3%) | 1 (1.1%) | 4 (25.0%) |
| Total | 16 (17.2%) | 17 (5.9%) | 16 (17.2%) |

**Fig 2.**
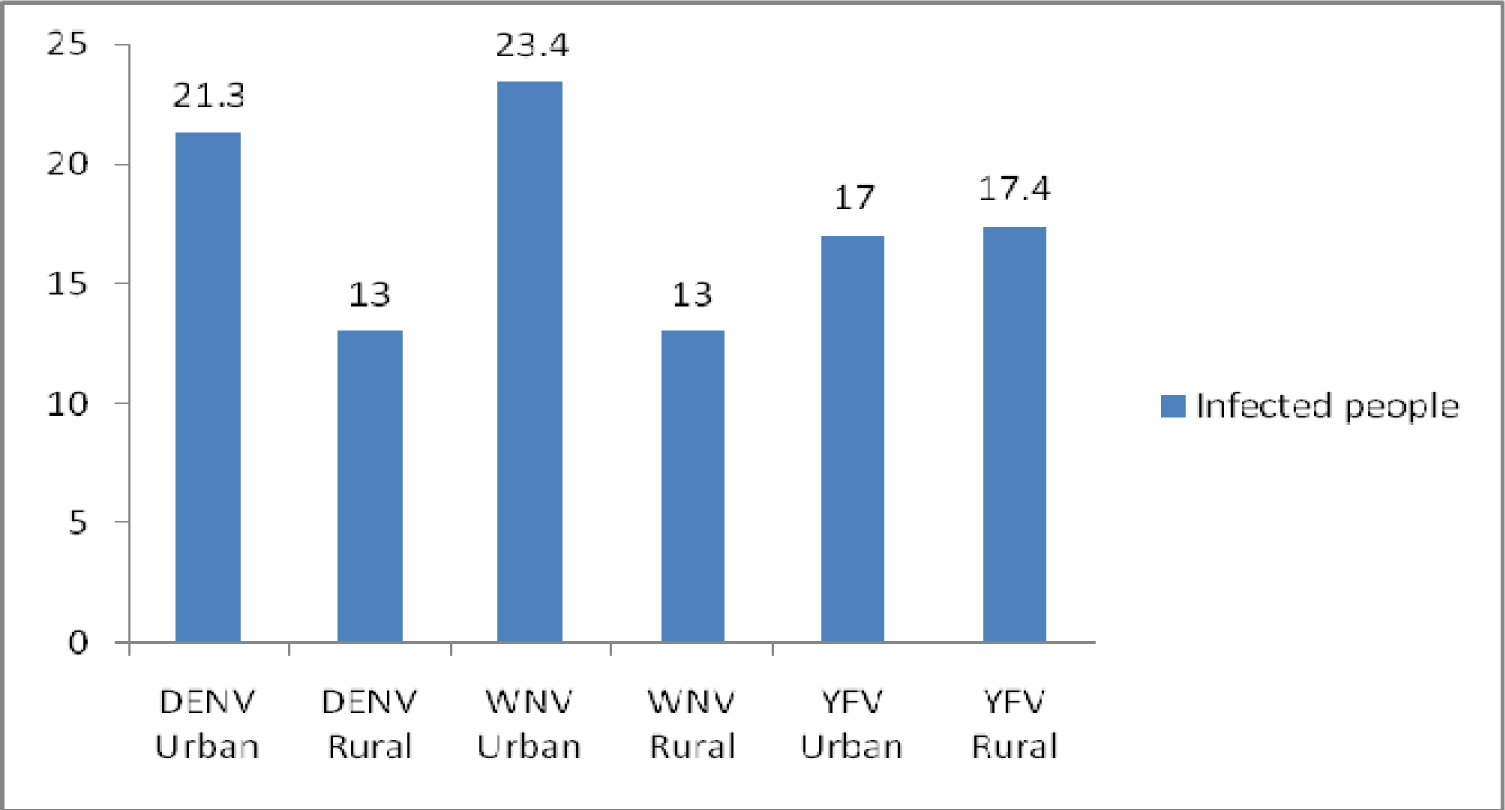
Distribution of proportion of infected people in rural and urban areas.

**Fig 3.**
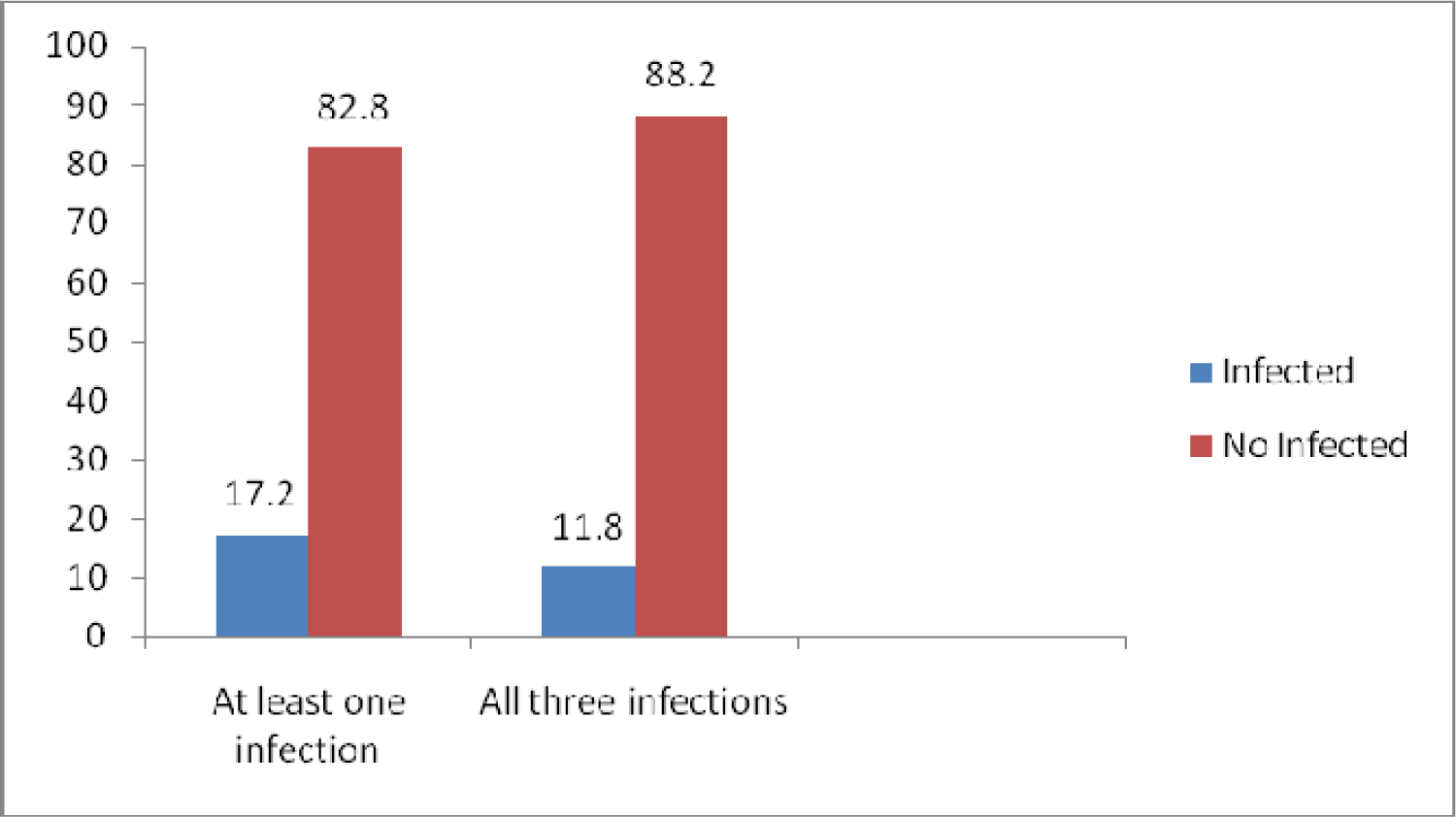
Distribution of individuals with dual or trio arboviral (DENV, WNV and YFV) infections.

## Discussion

Mosquitoes live with us in the habitat that we create. However, different land use establishes a range of habitats that can vary in quality and suitability for these mosquitoes. Human movement will continue to circulate vectors and pathogens. Throughout much of human evolutionary history, people have introduced exotic species into new geographic regions.

Epidemics of Flaviviruses have been documented throughout Africa over several decades. The spread of these viruses are limited by the presence of the mosquito vector which is found in tropical and subtropical climatic zones. Diseases caused by flaviviruses are the next most common tropical arthropod borne disease after malaria. This is more worrisome because of the presence of the vector in rural settings as well as urban cities (WHO, 2014).

The occurrence of arbovirus infections is an indication of the competence of mosquitoes. Although epidemics have not been identified in the study area, yellow fever epidemics occurred in Ogbomosho in 1946 and in Eastern Nigeria in 1951-53 (MacNamara, 1954). In 1987, there were epidemics of yellow fever in many parts of Southwestern Nigeria (Ogbomoso inclusive), which prompted mass vaccination of millions of Nigerians in the late 90s and stopped because of financial reasons but continued in 1997.

In sub-Saharan Africa, surveillance data for arboviral diseases is scanty, routine testing is not done and misdiagnosis is common because patients do not present with classical symptoms. In tropical countries there is increased hospitalization due to febrile illnesses which were initially suspected to be malaria or typhoid fever (Baba *et al.,* 2006).

In this study, an overall prevalence of 52.7% (49/93) from the recruited individuals. Out of the 49 (52.7%) IgM positive patients, 16 (17.2%) were positive for DENV, 17 (18.3%) for WNV and 16 (17.2%) for YFV. The prevalence recorded for DENV is higher than 0.5% prevalence recorded at Maiduguri (Baba and Talle, 2011). The result obtained is lower than the prevalence rates of 30.8% (Faneye *et* al., 2013), 35% and 73% (Oyero and Ayukekbong, 2014). The result is the same with the result obtained in Ogbomoso (Oladipo *et al*., 2014). The 18.3% prevalence obtained for WNV is lower than a 44% prevalence obtained by Fagbami (1977) at Igbo-Ora. The result (17.2%) obtained for YFV is lower than 36% prevalence obtained at Igbo-Ora (Fagbami, 1977). The disparity in the results might be due to differences in the specificity of the serological tests used, sample size and the period when the studies were carried out.

High prevalence were recorded in the age groups 16-30, 31-45 and 61-75 years in the age distribution of IgM positivity. This is in accordance with the findings of Oladipo *et al*., (2014) that reported high prevalence among age groups 16-30 and 31-45 years which are within the age range of this study.

Gender analysis of the positive samples showed higher prevalence among females than males. This is consistent with the previous observations in Mexico (Kaplan *et al*., 1983). However, it disagrees with the findings of Awando *et al*., (2013) and and Oladipo *et al*., (2014). The difference might be as a result of difference in sample size.

The result also shows that patients from the urban settings had more of the infections than those in the rural areas, however, for YNF, higher prevalence was found in the rural area. This is consistent with the findings of Collenberg *et al*., (2006) and Oladipo *et* al., (2014) that the prevalence of DENV was higher in urban settings than rural settings in Burkina Faso. This statement cannot be generalized because of the difference in sample size and testing methods.

The prevalence of dual and trio arboviral infection showed 17.2% and 11.8% respectively. Co-infections of arbovirus has been reported in Nigeria (Baba *et al*., 2012). Co-infections have been found to provide opportunity for genetic materials exchange and mutations resulting in the emergence of strains with enhanced disease severity (Ayukekbong, 2014).

In conclusion, the result obtained confirms the circulation of Flaviviruses in Ogbomoso. There should be proper diagnosis of these infections and public awareness on effective vector control.

